# Derivation of aerial insect concentration with triple frequency cloud radar observations

**DOI:** 10.1101/2024.07.30.605781

**Authors:** Moritz Lochmann, Birgen Haest, Teresa Vogl, Roel van Klink, Freya I. Addison, Maximilian Maahn, Willi Schimmel, Christian Wirth, Johannes Quaas, Heike Kalesse-Los

## Abstract

Aerial insects are vital for nature and society. Though methods to observe flying insects have consistently improved in the last decades, insects remain difficult to monitor systematically and consistently over large spatial and temporal scales. Remote sensing with radars has proved to be one of the more effective tools for observation. However, as radars are most sensitive to targets that are of similar size to the radar wavelength, the detectable sub-group of aerial insects of a certain size range depends on the employed radar. Here, we present a novel method based on spectral data of zenith-pointing Doppler cloud radars to estimate insect concentration in a vertical profile. Multiple meteorological state-of-the-art algorithms are combined to extract insect signals from the radar data and quantify their abundance from 150 m to 3000 m above the ground. For evaluation, this method is applied to triple frequency data from X-band (*λ* = 3.2 cm), Ka-band (*λ* = 0.85 cm) and W-band (*λ* = 0.32 cm) Doppler cloud radars from a three months summertime observation period in Germany. Observations of a case study of two elevated aerial insect layers are presented. Furthermore, differences in diel cycles of aerial insect concentration (*aic*) obtained from the three radar instruments are compared. With its superior sensitivity to very small insects like aphids, W-band shows the highest aerial insect concentrations during day-time in the lower altitudes. Generally, the higher wavelength radars show higher aerial insect concentrations in higher altitudes (Ka-band) and during the night (X-band).

## II. Introduction

OVER the past decades, multiple studies have reported strong declines in insect population sizes [1], abundances [2], [3] and biomass [4]. Most of these studies have relied on the standard trapping of insects and subsequent identification of the trapped individuals [5]. However, the data from such traps only represents the aggregated insect assemblage over a certain sampling period (usually a week or more), and for an unknown sampling area, and thus the potential to derive the spatio-temporal distribution of insect activity is limited. In addition, the processing of the trapped insects is extremely laborious and, hence, there is a growing need for non-invasive, low maintenance methods for measuring changes in insect abundance, biomass and movement [6].

A variety of methods have been introduced to automatically monitor insects during flight. Insects have been observed by video cameras [7], optical sensors [8] and lidar instruments [9], [10], but radar has proved to be one of the most effective tools for observing insect flight [11], [12]. As remote-sensing instrumentation, radars operate in a stand-alone way and on a time resolution usually ranging between seconds (cloud radars) to minutes (weather surveillance radars). In addition, zenith-pointing radars allow sampling of the entire vertical atmospheric column above the instrument. Scanning radars further have the potential to cover spatially large regions. During recent years, the deployment of specialized radar-based detection methods for aerial fauna has become more prevalent, especially for movements of up to several hundred meters up in the air [12]–[17]. To study larger flying animals, like birds and bats, radar instruments with wavelengths of a few up to several centimeters, like X-band (*λ* = 2.5-3.75 cm) [18], [19], C-Band (*λ* = 4-8 cm) [20], [21], S-Band (*λ* = 7.5-15 cm) [22], [23], and L-band (*λ* = 15-30 cm) [24] are ever-increasingly used. As radars are most sensitive to targets that are comparable in size to their wavelength, radar instruments operating at lower wavelengths are likely more useful for the detection of aerial insects [25]. Nevertheless, S-band, and more rarely also C-Band [26] weather radars have been shown useful to study high-density insect movements [27], including specific events like mayfly emergences [28], [29]. X-Band radars have repeatedly been shown useful to study larger individual insects [6], [12], [13], [16], [30], but mostly miss insects with smaller radar cross sections [31], which, in fact, are often more abundant [32]. Ku-Band (*λ* = 1.67-2.4 cm), K-Band (*λ* = 1.11-1.67 cm) and Ka-Band (*λ* = 0.75-1.11 cm) radars have seen some use for entomological purposes [15], [33]–[35], however, only few studies have been published on W-Band (*λ* = 0.32 cm) radar insect detection [36], [37].

Cloud radars have become established tools in atmospheric research, and their publicly available data present a valuable opportunity to study aerial insect activity. However, previously, entomological studies investigating cloud radar observations often either relied on using the radar reflectivity as a proxy for the number of insects present in each range gate [38] or used a Doppler spectrum width threshold to filter for single insect situations [33], [39] and, thus, did not consider situations when multiple insects occur in the same radar volume.

In the meteorological research community, multiple retrievals have been developed to distinguish different target particles detected by radars [36], [40]–[44], even when present in the same radar range gate. In this study, we explore how to effectively combine different established and state-of-the-art radar data processing techniques to achieve novel results in an aero-ecological context. We present a new aerial insect detection method based on Doppler cloud radar observations which we apply on triple frequency observations from W-, Ka- and X-band Doppler cloud radar instruments. We expect the W-Band observations to detect smaller insects in the size range of, e.g., aphids, while X-Band radar is more sensitive to larger insects, like moths [16], [45].

## III. Instruments and software tools

This section introduces the three radar instruments used in this study: the X-band radar MIRA10 (9.4 GHz, *λ* = 3.2 cm), the Ka-band radar MIRA36 (35.5 GHz, *λ* = 0.85 cm) and the W-band radar RPG94 (94 GHz, *λ* = 0.32 cm). In the following, the radar instruments will be referred to by their respective frequency band. Subsequently, the utilized software tools for flying insect detection from cloud radar Doppler spectra are presented individually. The algorithm chain of CloudnetPy (Sect. III-B) is used to create target classification labels for defining an insect mask. Afterwards, the radar Doppler peak detection algorithms PEAKO-peakTree (Sect. III-C) are used to extract information on insect quantities and properties. Section IV details how these software tools are applied to the radar data to detect insects.

### A. Doppler radar instruments

For the purpose of this study, triple frequency cloud radar observations from 1 June 2019 to 31 August 2019 at the Jülich Observatory for Cloud Evolution Core Facility (JOYCE-CF, [46], 50°54’ N, 6°24’ E; 111 m a.s.l., Central European Time Zone CET) west of Cologne, Germany, were evaluated. The setup of these instruments is similar to previous field experiments such as TRIPEx [47] and TRIPEx-pol [48], which focused on winter precipitation. The X- and Ka-band radars are pulsed radars manufactured by Metek [49], [50]. The Ka-band radar transmits linearly polarized pulses and then receives co- and cross polarized returns simultaneously. This enables the derivation of the linear depolarization ratio (LDR) which is the ratio of cross-to co-channel reflectivity and gives information about the target shapes [47]. Although LDR might be a valuable asset in detecting insects, this study does not explicitly require polarized cloud radar measurements to avoid excluding single-pol instruments. The W-Band Doppler cloud radar used in this study is a 94 GHz frequency modulated continuous wave (FMCW) non-polarimetric instrument developed for meteorological applications [44], [51]. For this FMCW radar, the transmitted frequency is modulated slightly (300–3600 kHz) around the center frequency of 94 GHz to continuously sample the atmosphere above the instrument in multiple layers, so-called chirps [51]. The user can set measurement parameters for each chirp individually in the chirp table configurator that is applied to a certain range. These chirps are observed consecutively. In addition to the two chirps relevant to our investigation, there are two more chirps above 3 km (Chirp 3: 3000 - 6000 m, 1.8 s integration time; Chirp 4: 6000 - 12000 m, 3.6 s integration time). Table I shows the most relevant measurement parameters for the data set analyzed here for the three radar instruments.

**TABLE I.** Radar settings for the three radar instruments. For the W-band radar, the settings for the first two chirps are shown.

| measurement parameter | X-band | Ka-band | W-band<br>chirp 1 | W-band<br>chirp 2 |
| --- | --- | --- | --- | --- |
| center frequency (GHz) | 9.4 | 35.5 | 94.0 | 94.0 |
| wavelength (cm) | 3.2 | 0.85 | 0.32 | 0.32 |
| temporal resolution (s) | 2.0 | 2.0 | 0.6 | 0.9 |
| considered range (km) | 0.14-3.0 | 0.22-3.0 | 0.15-0.8 | 0.8-3.0 |
| range resolution (m) | 36.0 | 36.0 | 29.8 | 11.9 |
| minimum detectable reflectivity at 500 m (dBZ) | -54 | -78 | -52 | - |
| Nyquist velocity ($\text{m s}^{-1}$) | 78.0 | 20.0 | 11.0 | 7.3 |
| Doppler velocity resolution ($\text{cm s}^{-1}$) | 3.8 | 4.0 | 2.15 | 1.42 |
| number of Doppler bins | 4096 | 1024 | 1024 | 1024 |
| half-power full beam width ($^{\circ}$) | 1.0 | 0.6 | 0.6 | 0.6 |

Figure 1a shows the data availability for the three radars during the three-month observation period. For W-band, there are two weeks of data missing in total during these three months, but for the other two frequencies there are almost no data gaps. Data availability is reduced for all instruments when we filter out profiles without insects (Fig. 1b). Out of 2208 possible hours of data in 92 operation days, 1717 (78 %) remain for W-band, 2008 (91 %) for Ka-band and 2063 (93 %) remain for X-band after applying the insect mask.

**Fig 1.**
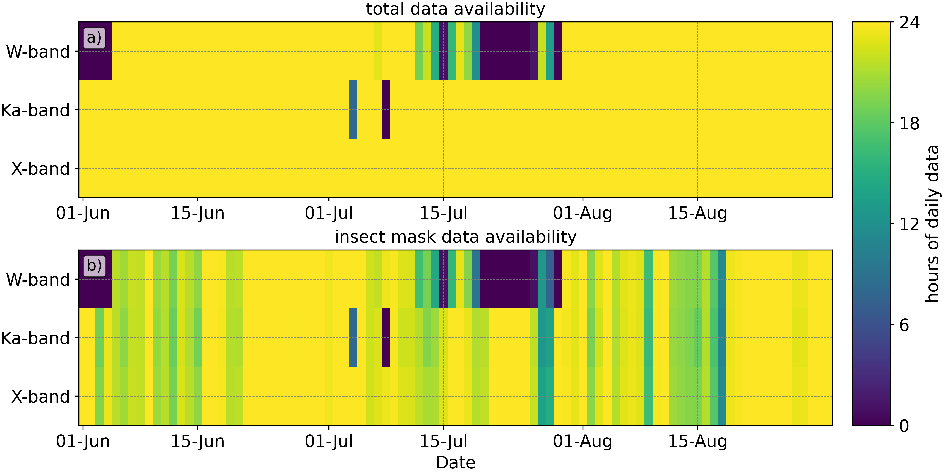
Daily data availability in hours of day for the three radars from June 1 to August 31, 2019. a) shows the overall data availability and b) shows the data availability after selecting only profiles containing insects.

The most prominent difference between the radar instruments is their wavelength (W-band: 0.32 cm, Ka-band: 0.85 cm, X-band: 3.2 cm) and, because of the associated sensitivity differences, the size ranges the radars are most sensitive to. For targets that are much smaller than the radar wavelength, the backscattering cross section is proportional to the 6th power of target size (known as Rayleigh scattering assumption). However, for targets similar to or larger than the radar wavelength the Rayleigh scattering assumption is no longer valid and instead scattering occurs in the Mie regime. In the Mie scattering regime, the backscattering cross section no longer scales with target size [52]. Instead, it oscillates with consecutive maxima and minima with increasing target size [53]. As a point of reference, for a conducting sphere with diameter D, W-Band scattering enters the Mie regime for approx. D>0.1 cm, Ka-Band scattering enters the Mie regime for approx. D>0.3 cm and X-Band scattering enters the Mie regime for approx. D>1 cm [11]. Thus, the larger the wavelength of the radar, the more reliably it is able to observe larger insects, e.g., moths, at the cost of not being sensitive to small insects in the mm-size range. With X-band, a larger size range of insects — ranging into the cm size range — can be detected in the Rayleigh regime. For increasingly smaller insects, the radar cross section’s inverse fourth-law dependence on wavelength is the limiting factor, making the use of higher frequency radar instruments preferable when aiming to observe very small targets [11], [50]. Thus, W-band more reliably observes smaller insects in the sub mm size range, like aphids. Another reason that the three instruments may not detect exactly the same targets is their sensitivity (see Tab. I). In our analysis, the sensitivity of the radar instruments is inferred from the first percentile of the detected reflectivity values. The Ka-band radar is most sensitive for the whole considered vertical profile. X-band and W-band radars are more comparable in the first W-band chirp up to 800 m (approx. 5 dBZ difference, not shown), but W-band signals are generally attenuated more strongly by water vapor.

All three instruments operate with a non-rotating vertically-observing radar beam with similar half-power beam widths (W-band, Ka-band: 0.6°, X-band: 1.0°). As an example, at 500 m altitude, the diameters of the observation volumes would be about 5 m for W-band and Ka-band and approximately 9 m for X-band, resulting in observation volumes of 641 m^3^ (W), 775 m^3^ (Ka), 2155 m^3^ (X) at 500 m altitude, respectively. Consequently, the X-band radar has the largest observation volume, which can be of advantage in situations with very low insect concentrations. During the integration time of the radar (see I) air is advected into the observation volume. This increases the volume of air sampled by the radar beyond the geometrically derived radar observation volume depending on the environmental wind speed *u*. Insects leaving or entering the observation volume during the integration time do not necessarily have the same radial velocity in regard to the radar instrument and, thus, appear at different positions in the radar Doppler spectrum. As the vertical wind component is usually much smaller than the horizontal component, only the effect of the latter will be considered. We calculate this additional wind-induced volume as the product of the amount of wind shift *u*·*t* over the trapezoidal area *A* of the radar observation volume perpendicular to the wind. The sum of the default observation volume *V*_0_ and the wind-induced volume yields the effective observation volume *V*_*EO*_ = *V*_0_ + *u* ·*t* ·*A*. Because *V*_0_ scales exponentially with altitude while the additional wind-induced volume (*u*· *t* ·*A*) only scales linearly with altitude, this correction decreases in impact with increasing altitude.

The steps for deriving the aerial insect concentration (*aic*) product from the radar data detailed in Sect. IV have to be applied to the raw spectral radar data. To enable a meaningful comparison afterwards, the data sets have been interpolated to the same time-height grid. We interpolated the output to a vertical resolution matching the W-band data, which has the finest resolution, and averaged to a temporal resolution of 5 s.

### B. CloudnetPy

The Cloudnet project is part of the Aerosol, Clouds and Trace Gases Research Infrastructure (ACTRIS) and provides a systematic characterization of atmospheric targets by analyzing continuous ground-based observations of the vertical profiles of the atmosphere [40]. Utilizing synergistic effects between Doppler cloud radar (for observation of hydrometeors), backscatter lidar (for observation of aerosol particles and liquid cloud base height) and microwave radiometer instruments (for atmospheric temperature and humidity profiling) in combination with atmospheric model data, Cloudnet is able to label the target particles at each point of the vertical profile into different classes. These classes include, e.g., ice, liquid, aerosol, insects and combinations of those. The JOYCE Cloudnet products are processed using data from the MIRA36 cloud radar, CHM15k ceilometer, HATPRO microwave radiometer and ECMWF Integrated Forecast System model data.

The CloudnetPy processing toolbox [54] provides a Python implementation of the full Cloudnet processing chain. The insect detection in CloudnetPy uses probability functions based on empirical thresholds of air temperature and radar variables. Cumulative probability distribution functions depending on the empirically chosen center values (mean value of the distribution) and scales (width of the distribution) are calculated for each of these threshold values.

There are two options for assessing the insect probability within CloudnetPy. The first option is based on the LDR. If the cloud radar is polarized and LDR is available, the variables used for determining the insect probability are LDR and wet bulb temperature, which is the lowest temperature to which air can be cooled by the evaporation of water in it, in the considered altitude from ancillary weather model data. The individual insect probability for both of these variables is calculated and multiplied to get the combined insect probability. This method is generally considered the best proxy for insect detection [54]. However, the LDR value is not always available, e.g., because of low signal to noise ratios or because some radar instruments are not capable of deriving LDR. In that case, the alternative variables used to calculate the insect probability are the integrated radar Doppler spectra data (moments) radar reflectivity, mean Doppler velocity, Doppler spectral width as well as wet bulb temperature. Both approaches largely depend on the empirically chosen threshold values for the radar variables. If the insect probability by either of these two approaches exceeds the CloudnetPy threshold of 0.8, the range gate is classified as ‘insect’. Please note that every non-hydrometeor Cloudnet pixel which exceeds this threshold is called ‘insect’. In reality, point targets other than insects, e.g., debris or plant material, might also fall into that category. Thus, we are aware that this approach is likely overestimating the true number of insects. Whether this effect can be quantified, and if it is especially prominent for high surface wind speeds, will be evaluated in a follow-up study.

### C. PEAKO-peakTree

If only one insect is present in a radar range bin, its signature in the Doppler spectrum can be easily identified, because it only causes one narrow peak. However, in summertime conditions, typically multiple insects are present in the radar observation volume, causing a superposition of multiple sharp narrow peaks in the Doppler spectrum [36], [42], as illustrated in Fig. 3. The supervised machine learning radar Doppler spectra peak-finding algorithm PEAKO [55] is used to investigate the characteristics of present peaks, even if they are not separated by the noise floor. Initially developed to determine the number of hydrometeor types (e.g., ice, drizzle or cloud droplets) in a certain cloud volume, PEAKO is trained on at least 200 radar Doppler spectra for X-band, Ka-band and each chirp section of the W-band radar. The training provides a set of parameters used to identify peaks in the typically noisy Doppler spectra for the specific cloud radar settings used. The six adjustable parameters are thresholds for number of time and range averages, the span and polynomial order for smoothing, peak width and peak prominence [56].

**Fig 2.**
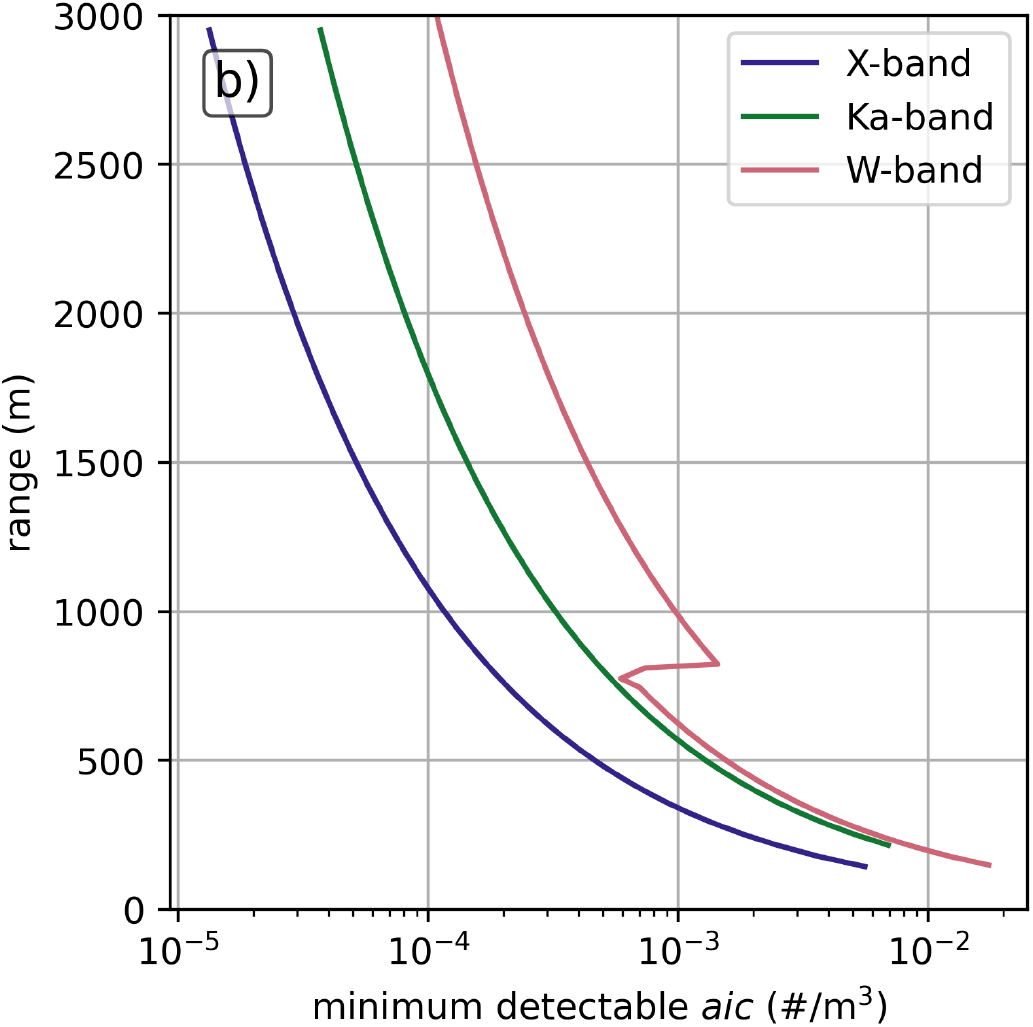
Theoretical minimum detectable aerial insect concentration without taking horizontal wind speed into account. The discontinuity in the W-band data is caused by the transition from first to second chirp (see Tab. I).

**Fig 3.**
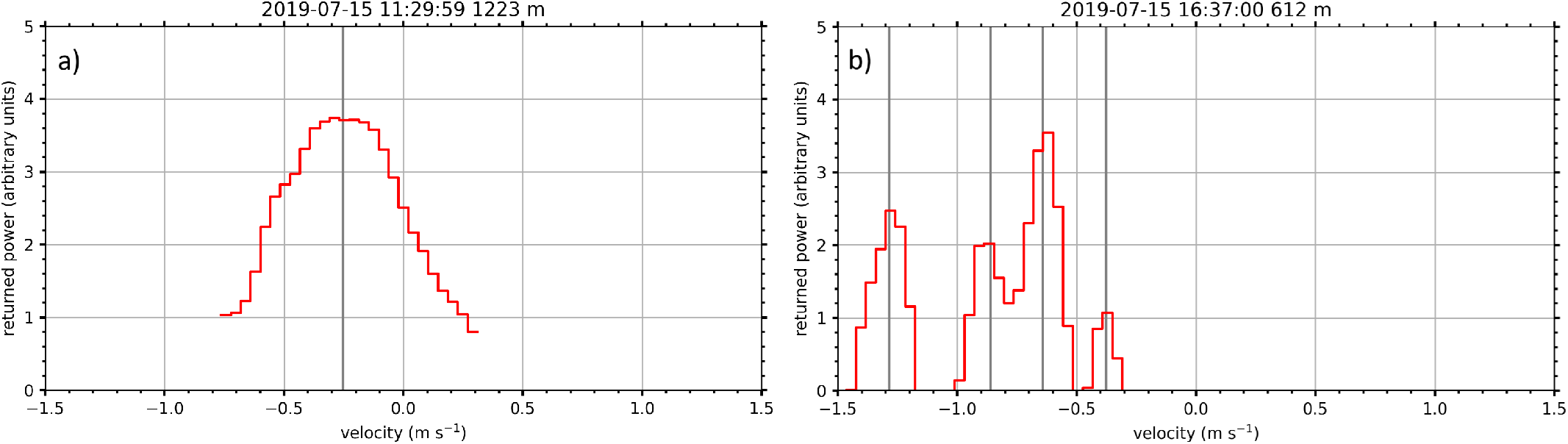
Comparison of typical cloud radar Doppler spectrum observing cloud droplets (left) and insect echoes (right) as obtained from Ka-band radar on July 15, 2019 at JOYCE. The presence of cloud droplets in the left case is substantiated by supplementary measurements from a ceilometer. The cloud droplets appear as a broader peak close to 0 m s^−1^, while the insects appear as individual sharp peaks at various Doppler velocities. Negative Doppler velocity values indicate a motion toward the radar (downward) and positive values a motion away from the radar (upward). The grey vertical lines represent Doppler spectrum peaks found by peakTree (see Sect. III-C).

For each of those parameters a range of realistic values are defined for the training of PEAKO. To identify insect peaks, we omitted the height averaging, because we expect each insect to always only be in one range gate. Although single flying insects may be detected in the same range gate for consecutive time steps, they are not expected to consistently do so. Early tests with temporal averaging over multiple radar time steps showed poor results and required significantly more training time, thus, temporal averaging was omitted for later training runs. Next, the radar Doppler spectra are smoothed according to span and polynomial order. The smoothing algorithm uses a certain window size (span), over which a polynomial is fit to the segment of the spectrum. The span was varied between 0.05 m s^−1^ and 0.3 m s^−1^ and the polynomial order between 2 and 3. The best-performing span value varied between the instruments. For all instruments, a polynomial order of 2 achieved best training results. Only peaks with a width larger than the PEAKO width threshold are considered [56]. Since we expect insect peaks to be very narrow, we tested threshold values between 0 m s^−1^ and 0.1 m s^−1^ and for all trainings threshold values just above 0 performed best. The peak prominence describes the power difference between the maximum of a specific peak and the minimum between it and the closest higher peak. For this parameter, we tested values between 0 dB and 3 dB. The prominence value leading to best performance results varied between the different radar instruments. Finally, the combination of parameters leading to the highest similarity with the user-defined training data set are saved in a configuration file and can be used as input for peakTree. More details of the PEAKO algorithm are introduced and demonstrated in [55] and [56].

Subsequently, the peakTree algorithm is used to identify, organize and interpret the peaks in radar Doppler spectra [41]. It provides a recursive list of all subpeaks in the spectrum without a priori assumptions on their structure. These subpeaks are then characterized by their mean Doppler velocity, reflectivity, spectral width, skewness, prominence and LDR. peakTree can directly utilize the PEAKO training parameters via a configuration file [56]. Although peakTree inherently provides the option to select specific hydrometeor types with threshold based rules, it does not differentiate between hydrometeors and biological targets, such as insects.

## IV. Methodology - radar data processing

This section shows how we apply the different established radar processing techniques introduced in detail in the previous sections to data from the three radar instruments. Figure 4 depicts the sequence of these methods as applied in this work. First, CloudnetPy generates the target classification product based on the cloud radar moments (reflectivity, mean Doppler velocity and spectral width), weather model data and lidar data. The target labels are used to derive the insect mask, which is later applied to the peakTree output data. However, a review of the extensive data set has shown that the CloudnetPy target classification sometimes gives unrealistic results at the edges of clouds and precipitation, where pixels are erroneously labeled as insects [57]. It is unclear whether this is due to partial beam filling or uncharacteristically low reflectivity values for liquid targets, but these obvious misclassifications are removed here. Specifically, the applied filtering algorithm removes pixels containing insects which directly border pixels containing drizzle/rain in either the time or range dimension (orange, see Fig. 4). From the location of the thereby selected pixels, two bins in each direction of the two dimensions, i.e., two range gates up and down and two time steps (about 1 min) before and after, are removed and, thus, not considered for the subsequent analysis. From the remaining range gates classified as insects, an insect mask (orange, see Fig. 4) is created. We are aware that this approach may remove a few valid insect pixels, however, we choose to preferably avoid false positives.

**Fig 4.**
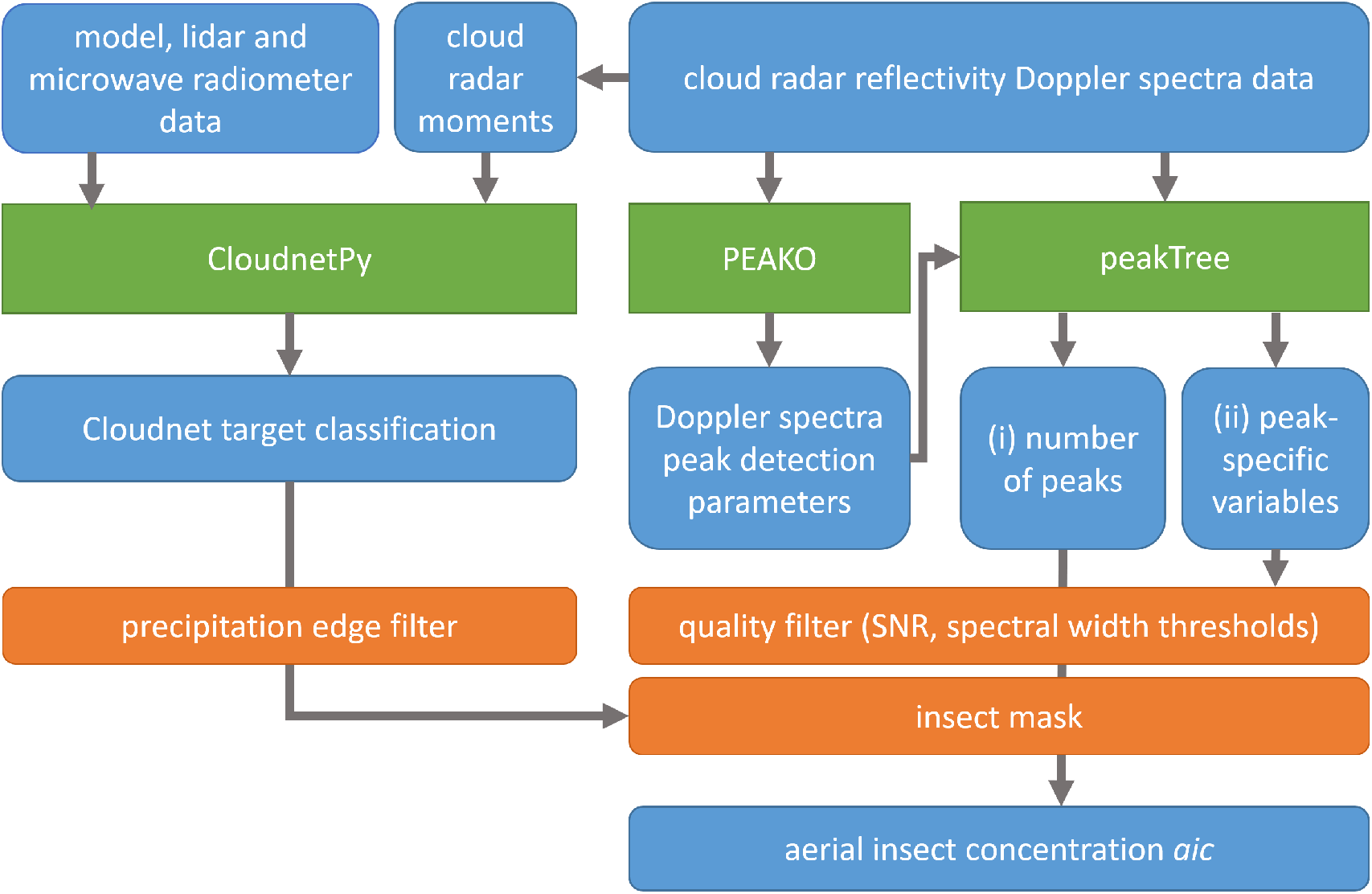
Data processing flow chart for aerial insect detection. Blue boxes represent used data sets, green boxes depict software tools and orange boxes represent filters which are applied on the data set.

To extract information about the number of insects in each range gate, the Doppler spectra are analysed with PEAKO and peakTree. It is assumed that every narrow and sharp peak in the radar Doppler spectrum of each ‘insect’ pixel represents one insect as established in [39]. Subsequently, the optimal parameters for the peak detection found by PEAKO are used to apply peakTree on the remaining data set. Finally, peakTree yields (i) the number and location of peaks in the radar Doppler spectra and (ii) the moments of the sub-peaks (reflectivity, velocity, spectral width, LDR) and other properties such as the prominence and signal-to-noise.

To increase the reliability of the detected peaks being insects, the spectral width and signal to noise ratio are combined in a reliability filter. Several different threshold values for these filters have been applied in a sensitivity study which will be summarized briefly.

The spectral signal to noise ratio (SNR) is calculated as the ratio between the spectral reflectivity and the mean noise level at each radar range gate. To determine the SNR of each peak, the spectral SNR is integrated between the lower and upper bound of each peak and then converted to dB. For the SNR, thresholds of 2 dB, 5 dB and 10 dB were evaluated. A SNR threshold of 10 dB is commonly used for hydrometeor populations (e.g., [48], [58]). For non-hydrometeor targets, [33] use a SNR threshold of 0 dB to filter for single insects. In our sensitivity study, we conclude that a more strict SNR threshold value of 5 dB retains the majority of the narrow Doppler spectra peaks.

The spectral width of each peak is defined as the width of the peak of the Doppler spectrum and calculated as the product of Doppler velocity resolution and the number of Doppler bins the peak spans across. Wood et al. [39] and Wainwright et al. [33] suggest using 0.1 m s^−1^ as maximum spectral width to assume that each peak is caused by a single insect. To investigate the sensitivity to this threshold, values of 0.05, 0.1 and 0.2 m s^−1^ are evaluated. More than 92-98 % of all peaks in insect pixels (depending on the radar frequency) have a spectral width lower than 0.1 m s^−1^. As this value seems a reasonable threshold it is applied for this study as well.

To ensure that the derived aerial insect concentration (*aic*) values are comparable between instruments, the number of detected insect peaks at each altitude is divided by the altitude-dependent effective observation volume. With this approach, the minimum detectable *aic* of individual measurements, defined as observing a single insect per range bin, decreases with altitude (see Fig. 2). Nevertheless, for the time-averaged analyses in the next chapter, *aic* values below this minimum value can occur when no insects are detected during some time steps of the averaging period. In addition, there is a maximum detectable *aic*, since we limit the number of peaks that peakTree may detect to 50. In practice, however, maximum *aic* values are at least a factor of 2 below this theoretical upper limit.

All three radar instruments observe the atmosphere with very high temporal resolution in the range of seconds. To improve the data consistency of all three radars and to reduce noise, the data is averaged temporally as explained in the following chapter.

## V. First aerial insect concentration results

To briefly demonstrate the potential of using Doppler cloud radar data for insect detection, the derived aerial insect concentration *aic* (in the following subscripted with the frequency band X, Ka or W) during the three month observation period is shown on different temporal scales. As explained in Sect. III-A, we expect the radar instruments to observe slightly different insect populations due to the wavelength and sensitivity differences. This assumption will be explored in this section. However, as this study is mainly supposed to introduce this method, more in-depth analyses will be showcased in future publications.

Typically, a clear dependence of *aic* on the evolution of the atmospheric boundary layer is evident for all considered radar frequencies. As cloud radar signals are able to penetrate clouds, they could theoretically detect flying insects above shallow cloud layers. In practice, however, this does not occur regularly. Figure 5a shows the time-height plot of a typical fair weather day (24 August 2019). The orange vertical lines represent times of sunrise at 5:35 CET and sunset at 19:36 CET. Insect activity begins to increase after sunrise. The maximum of diel insect flight at most altitudes is reached in the afternoon, approximately between 13 CET and 16 CET. Although radar sensitivity decreases with altitude, on some days we can detect insects in altitudes of up to 4 km with the highly sensitive Ka-band radar. After sunset, both the value of *aic* in most altitudes as well as the highest reached altitudes decrease for all wavelengths. During the night, *aic* values are typically lower than during the day. These diel patterns are in line with previous observations made with radars [16], [59].

**Fig 5.**
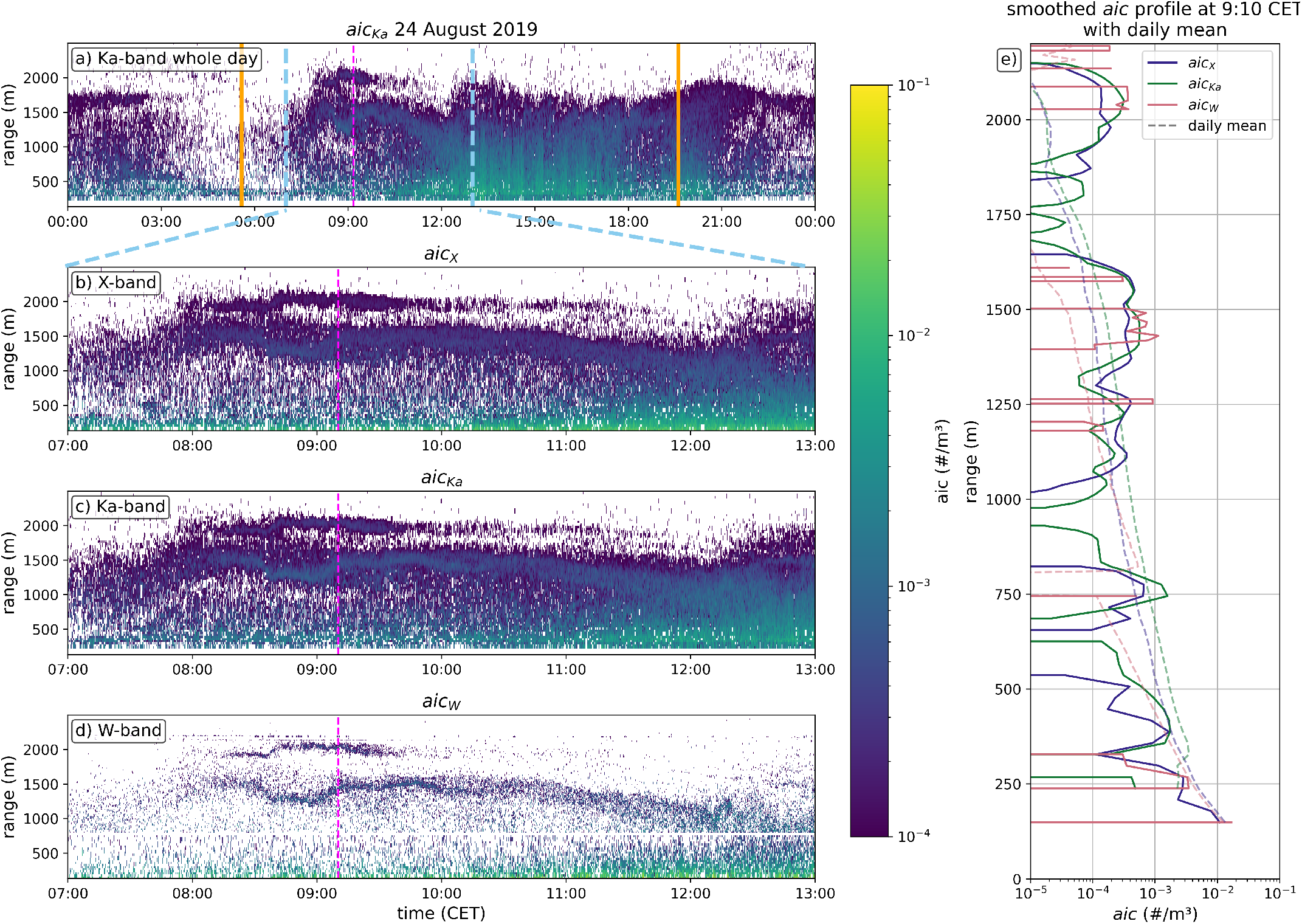
Case study of two elevated insect layers on 24 August 2019. The time-height plot a) shows *aic*_*Ka*_ for the whole day (note the different x-axis range). Orange vertical lines indicate sunrise and sunset. The light blue vertical lines indicate the time period shown in b)-d). The pink line indicates the time of the profile shown in e). The time-height plots b)-d) show *aic* for the three radars for the six hour time period between 7 CET and 13 CET. e) shows the smoothed *aic* profile (5 min mean centered around 9:10 CET) as solid, colored lines for the different wavelengths. The dotted lines represent the daily mean *aic* profiles.

We focus on a six hour time period (7 CET to 13 CET), where two elevated insect layers are detected by the cloud radars. Figure 5b-d show *aic* time-height plots for all three wavelengths up to an altitude of 2500 m. There are two features present, which are superimposed on the typical diel cycle of insect activity. One shallow elevated layer of insects occurs between approximately 8 CET and 10 CET centered at 2000 m above ground. Another, more pronounced layer is evident at approximately 1500 m altitude. This second layer also starts around 8 CET but continues until it merges with the typical day-time *aic* increase around noon (see also Fig. 5a). Although the W-band radar detects less insect echoes at these altitudes than the other two instruments, it is still able to represent these features. Figure 5e shows the vertical profile of *aic* at 9:10 CET (indicated with a dashed vertical pink line in Fig. 5a-d). The solid lines in e) are smoothed vertical profiles showing the mean *aic* in a 5 min time window around 9:10 CET, the dashed lines are daily mean values for each altitude. Generally, at this time of day, the convective atmospheric boundary layer is not fully developed yet. Below the two elevated layers, the *aic* profiles (solid lines) are consistently below the daily mean *aic* profile. However, at the altitude ranges of these two layers (1300-1600 m and 1900-2100 m) the current *aic* profiles show higher values than the daily mean.

This example showcases the high spatial and temporal resolutions with which *aic* can be resolved with this newly developed method. However, looking at longer time periods and condensing the information can enable generalizations. To smooth the dataset, the *aic* data is resampled to 10 min time periods. A 10 minute interval was chosen as a temporal grid to compromise between having a smooth data set and still being able to exploit the high temporal resolution of the radar instruments in representing changing diel insect activity patterns. Especially the very high temporal resolution remains a valuable asset for the investigation of even small scale insect activity patterns. The mean number of simultaneously observed peaks was chosen to represent each time interval. Note that when the radar does not detect any insects, the range gates in question are filled with zeros which are taken into account when averaging. Time periods for which the radar data is masked due to occurring hydrometeors or data gaps are filled with NaN values and are thus not taken into account when averaging.

Figure 6 shows the mean diel cycle of the absolute *aic* values divided into monthly subplots. In addition to the highest *aic* values occurring around midday, maxima around dawn and dusk are visible for all wavelengths, although with different intensity. This diel cycle qualitatively compares well to previous findings of, e.g., [16], [45], [60] which all use X-band radar observations. The shaded standard deviation in Fig. 6 shows that the variability in the derived *aic* value sometimes reaches one order of magnitude, but differences between the three wavelengths are still apparent. When comparing the diel cycle of the mean *aic* from August (bottom plot in Fig. 6) to the other months, it is evident that *aic* maxima at dawn and dusk occur later in the morning (around 5:30 CET, compared to 4:00 CET in June and July) and earlier in the evening (around 19:00 CET, compared to 21:00 CET in June and July), likely related to the shorter daylight period in August compared to June and July. In addition, mean *aic* values throughout the day are higher in June than in August. Variations in *aic*_*X*_ during the day are weaker than for the other two wavelengths. In July and August night-time *aic*_*X*_ is higher than *aic*_*Ka*_ and *aic*_*W*_, which is indeed to be expected if mostly larger insects are flying during the night [16].

**Fig 6.**
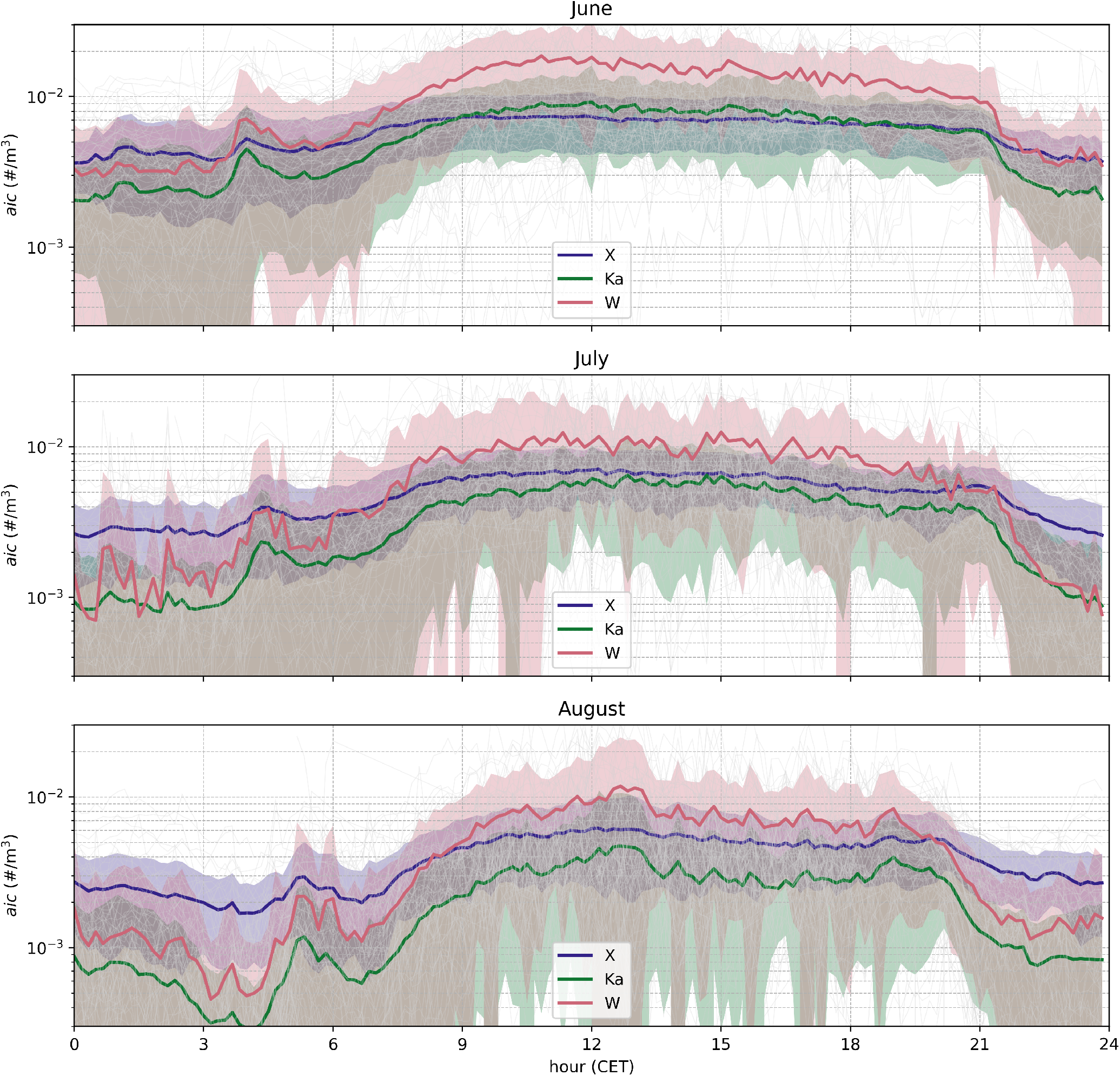
Mean diel cycle of absolute *aic* values in 250 m. Thin grey lines represent the diel cycle of each day. The solid colored lines represent the mean diel cycles for the different frequencies. Shaded areas represent one standard deviation for each wavelength’s mean value.

Because radar sensitivity to targets of certain sizes depends on wavelength, the shorter-wavelength radar (e.g., W-Band) can detect smaller insects that the longer-wavelength radar (e.g., X-Band) cannot, given similar instrument sensitivities. If the *aic* values derived from both radars in a certain range gate do not change equally over time, we can thus conclude that this is caused by insects only one radar is sensitive to. To investigate this, the *aic* of X-band and W-band (the two radars with the largest wavelength difference) are compared: if *aic*_*W*_ at, e.g., 500 m altitude decreases at night, but *aic*_*X*_ remains constant, we can conclude that we observe less small flying insects that only the W-band is sensitive to. The *aic* ratio *aic*_*X/W*_ = *aic*_*X*_*/aic*_*W*_ is presented in Fig. 7. If *aic*_*X/W*_ > 1 (*aic*_*X/W*_ < 1), *aic*_*X*_ is higher (lower) than *aic*_*W*_, indicating less (more) smaller insects, respectively.

**Fig 7.**
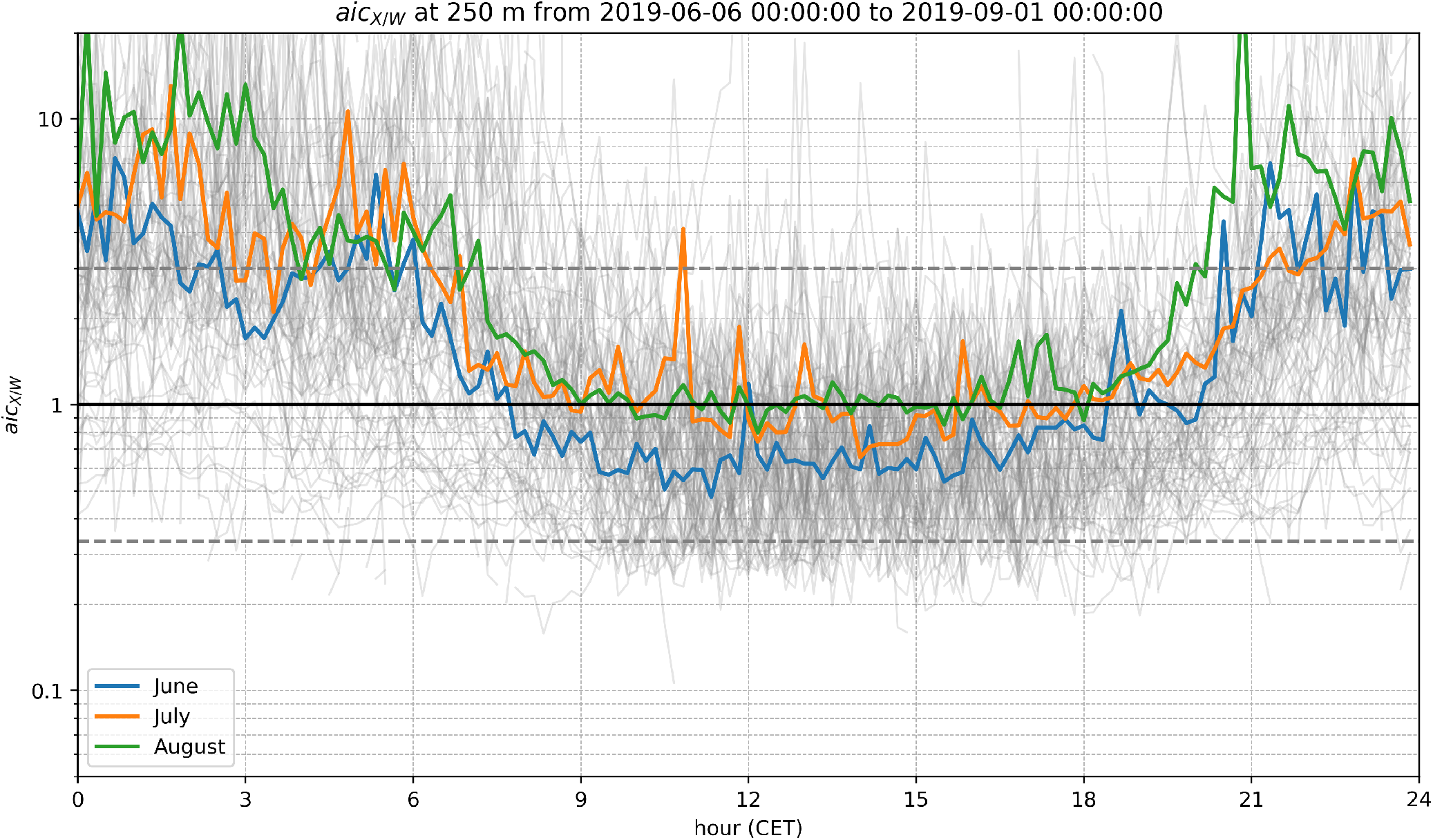
Mean diel cycle of *aicX/W* ratio in 250 m. Thin grey lines represent the diel cycle of each day. The solid colored lines represent the mean diel cycles for each month of the observation period (June, July, August). The horizontal solid line represents 1, when *aic*_*X*_ = *aic*_*W*_ . The horizontal dashed grey lines represent a factor of 3 in each direction.

We analyzed 10 min *aic*_*X/W*_ values for the three month observation period in Fig. 7. To include information for a longer time period the *aic* values for each daily 10 min interval are averaged monthly, that means, e.g., the *aic* values for every 0:00-0:10 CET window in June are averaged. The mean diel cycle of *aic*_*X/W*_ is shown as the colored lines in Fig. 7. As the time of sunrise and sunset - which presumably affects insect flight behaviour - change considerably during three months, we derived the mean diel cycle of *aic*_*X/W*_ for each month individually. A diel cycle of *aic*_*X/W*_ is evident in Fig. 7. During the night, values are highest and consistently above 1, meaning *aic*_*X*_ > *aic*_*W*_ . During the day, *aic*_*X/W*_ decreases to values below 1 (June, July), which means more insects were detected with the W-band radar than with the X-band radar. Thus, according to the assumption above, from these results we expect less smaller aerial insects during the night (high *aic*_*X/W*_ ratio) and more smaller aerial insects during the day (lower *aic*_*X/W*_). In August, *aic*_*X/W*_ decreases only to values of around 1 during the day. This could possibly be related to less small aerial insects being active later in summer. In the evening, the *aic*_*X/W*_ values increase toward the night-time maximum.

Lastly, we investigated day-to-day differences between absolute *aic* values for the three frequencies during the three month period. To keep this analysis simple, we choose the daily maximum value of *aic* in different altitudes if at least 18 hours of data are available for that day. Although Ka-band measurements start only at 215 m, the lowest altitude included in this overview is 150 m. Over the course of the data set, no obvious trend is visible. Therefore the results are presented as histograms (Fig. 8) for visual clarity. Generally, the maximum *aic* value decreases with altitude for all considered frequencies. For all altitudes, the maximum *aic*_*X*_ values are generally lower than for the other two frequencies. In the lowest range gate (150 m), maximum *aic*_*W*_ values are about three times higher than the maximum *aic*_*X*_ values. Also at 250 m, *aic*_*W*_ has the highest maximum values on most days, followed by *aic*_*Ka*_ and *aic*_*X*_ . At 500 m and 750 m altitude, maximum *aic*_*Ka*_ values are about twice as high as *aic*_*X*_, while maximum *aic*_*W*_ has a high variability ranging between *aic*_*X*_ and *aic*_*Ka*_.

**Fig 8.**
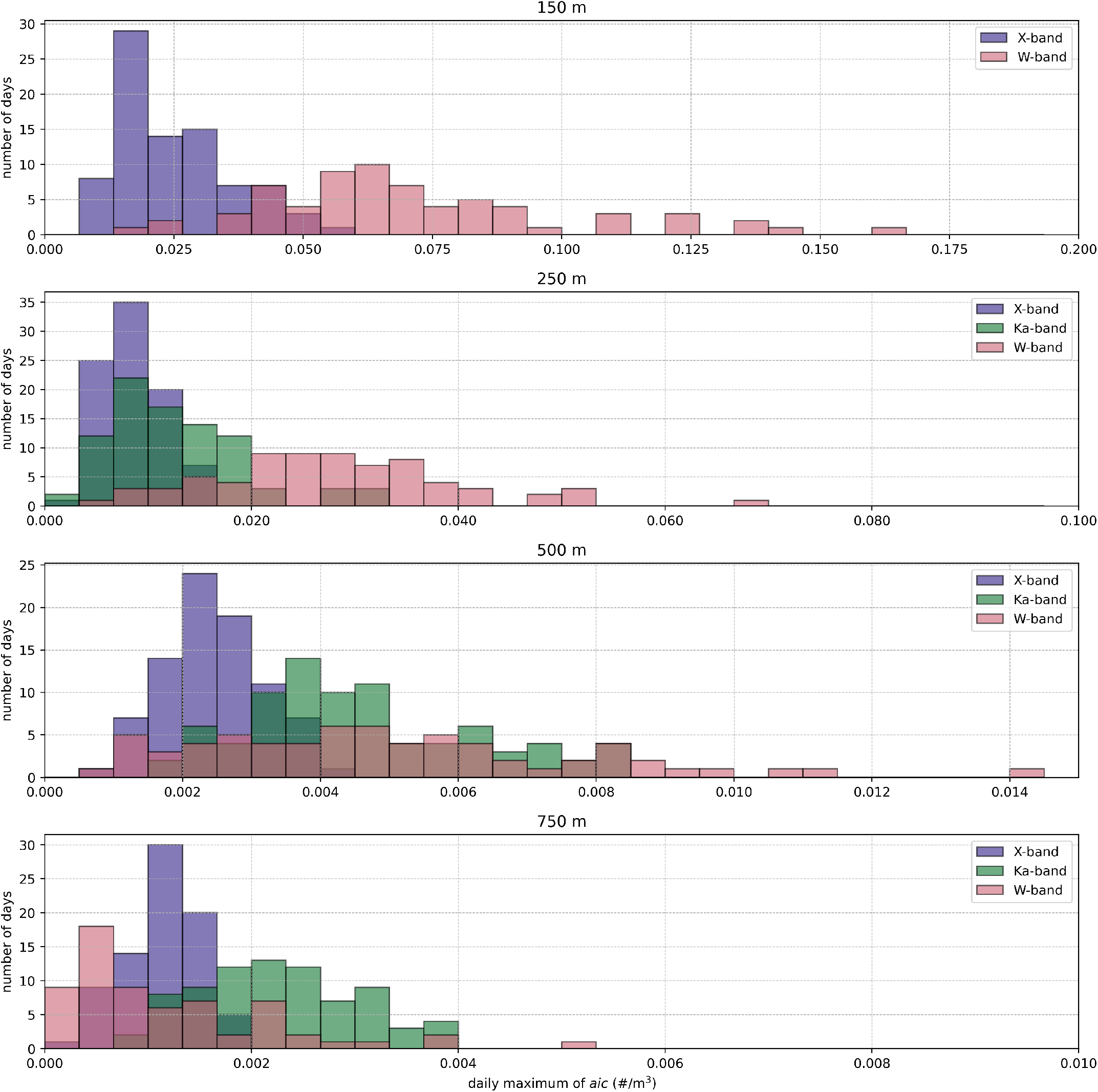
Histograms of maximum daily *aic* in 150 m, 250 m, 500 m and 750 m derived from the three radars. Note the different x-axis ranges.

Other studies investigating insect concentration values above ground include [60] (X-band) and [61] (aerial netting). More recent ones also present some vertical profiles from Ku-band [34], [62] and X-band radars [63]. However, please note that these literature values are mostly based on case studies instead of large datasets like ours and are taken in different regions of the world (Australia, Mongolia, England). The *aic* values found in our study are higher than those found in the mentioned studies. For example, at an altitude of 250 m above ground, values for insect concentration reported by other studies range between 5.0 ·10^-5^ m^−3^ [34] to 1.3 ·10^-3^ m^−3^ [62], while we found *aic* values ranging between 2.0 ·10^-3^ m^−3^ to 2 ·10^-2^ m^−3^ (monthly means) given in Fig. 6. However, as mentioned previously, standard deviation for our *aic* can reach one order of magnitude. There are several possible reasons why our absolute *aic* values differ from previous studies. Many estimates so far are based on data from dedicated biological radars [16], [60], [63], which report having low sensitivities to smaller insects [17]. Additionally, other radar specifications - like minimum detectable reflectivity - vary significantly between studies or are often not clearly reported. Furthermore, the *aic* values depend strongly on landscape type of the site [45], season, climate zones and probably exhibit strong interannual variability, which could explain differences between our values and previously reported values.

In our dataset, the lowest altitude above ground for which we derive *aic* is 150 m. To observe profiles of flying insects with remote-sensing techniques below that altitude, lidars can be used. Most lidar-based insect studies use the Scheimpflug lidar method and typically observe insects from few meters above the ground up to altitudes of approximately 100-150 m (e.g., [10], [64]). Scheimpflug lidars thus can be considered as complementary to radar in synergistic remote-sensing techniques for observing flying insects.

## VI. Summary, conclusions and outlook

This paper discussed the combination of several state-of-the-art radar data processing techniques to achieve novel results in the detection of flying insects using cloud radars operating at three different frequencies, which are sensitive to different size ranges of insects. The Cloudnet target classification was used to infer time-height pixels containing insects. Subsequently, the peak-finding algorithms PEAKO and peakTree were used to derive the number of insects present at each radar range gate. Note that this approach does not use peak signal power to infer target properties. Instead, a peak in the radar Doppler spectrum is taken only as evidence of a flying insect’s presence, regardless of its size, shape or species.

With the assumption that every narrow Doppler spectrum peak is caused by a single insect, *aic* values were calculated for X-band, Ka-band and W-band Doppler cloud radars at the Jülich ObservatorY for Cloud Evolution (JOYCE) for summer months 2019 (June, July, August). Given that radars are most sensitive to targets comparable in size to their wavelength, it is presumed that these instruments observe slightly different groups of aerial insect fauna. Differences in *aic* values derived from the radars, thus, help to identify and compare flight behaviour between smaller mm-size and larger cm-size insects. Knowledge about the flight timing and flight altitude of small insects derived from cloud radar data expands and fills in certain data gaps with additional information on a different subset of aerial fauna.

We demonstrated the potential of this novel insect detection method on a case study of two elevated insect layers detected between 1300 m and 2100 m. Although these features are detectable with all three radar instruments, they are less pronounced in the W-band data, which can be attributed to the lower radar sensitivity at this altitude. Comparing the vertical *aic* profile at the time of the features with the mean *aic* profile shows a significant increase in *aic* at the features altitude compared to the daily mean.

Afterwards, the mean diel cycle of the ratio *aic*_*X/W*_ was presented. From the wavelength dependency of the radars sensitivity we assume that the ratio *aic*_*X/W*_ changes when the size distribution of detected insects changes. Although we intentionally do not derive insect sizes from the cloud radar Doppler spectra, some information about the insect sizes can be gained from this evaluation. From this we concluded that small flying insects are less numerous during the night, which agrees with current entomological research.

Lastly, the maximum daily *aic* were evaluated in three altitudes. In general, *aic*_*X*_ values are lower than *aic*_*Ka*_ and *aic*_*W*_ . In 250 m altitude, maximum *aic*_*W*_ are often higher than maximum *aic*_*Ka*_ values. In the highest considered altitude 500 m, *aic*_*Ka*_ values are often highest. This can be attributed to the Ka-band radar having the best sensitivity (see Tab. I).

This study introduced the methodology of deriving *aic* values from triple frequency spectral Doppler cloud radar data and the application on a three months data set. The main focus of this publication is on introducing this methodology. A case study and several statistical results were presented to highlight the potential of the data set. Especially the difference in derived *aic* from different wavelengths promises novel results in an aeroecological context.

As we are able to derive aerial insect concentration (*aic*) from data, which were initially gathered for meteorological purposes, there are no additional costs for instrument acquisition, setup or maintenance associated with this approach. It should be noted, however, that the internal signal processing of these instruments is designed to most effectively extract the relevant meteorological properties. In this study, we had no means to modify these internal processes, but ways of adjusting them could potentially enhance the detection of insects from meteorological radar data. For beam-matched radars that see the same observation volume simultaneously with two frequencies (e.g., RPG 35/94 GHz dual frequency Doppler cloud radar), this approach could even be extended to derive characteristic insect features for every peak.

In a follow-up study we will apply this approach to multi-year radar observations performed at Cloudnet sites in different climate zones. Using larger data sets presents an opportunity to evaluate seasonal cycles as well as interannual trends and the potential dependence of *aic* and diel flight timing on environmental factors, e.g., land surface type, surface wind speed or air temperature. Applying the methods presented in this study to multi-year data sets may provide an opportunity to study these trends in aerial insect behaviour with high temporal and vertical resolution.

## Acknowledgments

This work was funded by the Saxon State Ministry for Science, Culture and Tourism (SMWK) – [3-7304/44/4-2023/8846]. RvK was funded by the DFG (FZT 118). We thank Stefan Kneifel (Ludwig Maximilian Universität Munich) and Bernhard Pospichal (University of Cologne) for providing the radar data set from JOYCE and Leonie von Terzi (Ludwig Maximilian Universität Munich) for providing help with the data set. We also thank Martin Radenz (Leibniz Institute for Tropospheric Research) for his work on peakTree and his help in the analysis of peakTree results. We acknowledge ACTRIS and Finnish Meteorological Institute for providing the Cloudnet data set which is available for download from https://cloudnet.fmi.fi. We acknowledge ECMWF for providing IFS model data.

## Contributions by the authors

ML was the primary author of this paper, did most of the data analysis and prepared the manuscript. BH provided important help in interpreting entomological results. TV helped greatly in setting up the PEAKO and peakTree training of the data set and provided help in interpreting results. RvK helped ML with the entomological analysis of the results. FA and MM helped in interpreting results and gave suggestions for the preparation of the manuscript. WS helped in the first stages of the manuscript with insect classification. JQ and CW helped in defining research questions and provided general guidance during the project. HKL developed the conceptual idea of this manuscript and helped greatly in interpreting results and preparing the manuscript.

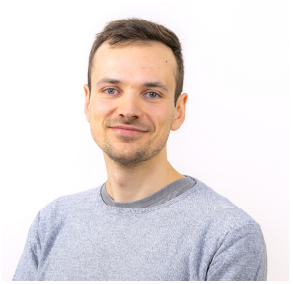

**Moritz Lochmann** received his B.Sc. and M.Sc. degrees in Meteorology at Leipzig university, Leipzig, Germany, in 2015 and 2018, respectively. Subsequently, he conducted his Ph.D. research in meteorology at the Leipzig Institute for Meteorology working on very short-term wind power predictions and received his degree in 2023. His current research centers on observing aerial insects with meteorological instruments, e.g. Doppler cloud radars.

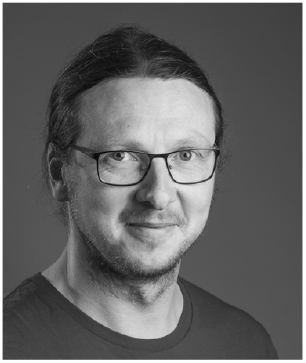

**Birgen Haest** received the M.Sc. in Bioscience Engineering degree from the University of Ghent, Belgium, in 2007. From 2008 to 2014, he did research on vegetation biodiversity mapping using optical remote sensing imagery at the Remote Sensing Department of the Flemish Institute of Technological Research, Mol, Belgium. In 2015, he moved to the Institute of Avian Research “Vogelwarte Helgoland” in Germany where, in 2019, he received his “Summa Cum Laude” PhD degree in ecology from the University of Oldenburg, Germany. Since 2019, he has been working as a Research Associate at the Swiss Ornithological Institute in Sempach, Switzerland, where he researches the migratory movements of birds, bats, and insects, amongst other using radar technology. In 2024, he was awarded a Mid-Career Award at the International Radar Aeroecology Conference, for his scientific contributions to the field of radar aeroecology.

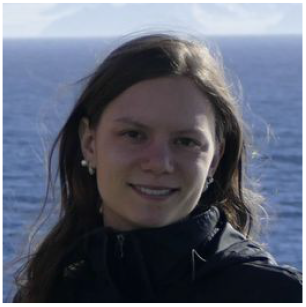

**Teresa Vogl** did her B.Sc. in environmental sciences at University of Bayreuth, Bayreuth, Germany, and her M.Sc. in meteorology at Leipzig university. Currently, she is doing her PhD on microphysical growth processes in mixed-phase clouds using cloud radar observations. Furthermore, she is also working on machine learning methodologies for photovolatics power prediction based on satellite observations.

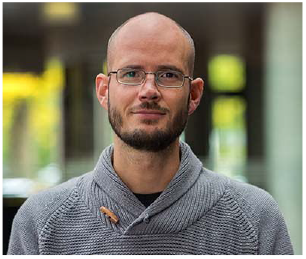

**Roel van Klink** is an entomologist that specializes on changes in insect biodiversity over time. His work has shown that declines in insect biomass and abundance are common, but not universal. Previously he worked on hands on conservation management for insects and on rewilding. He is also an avid collector of plant- and leafhoppers.

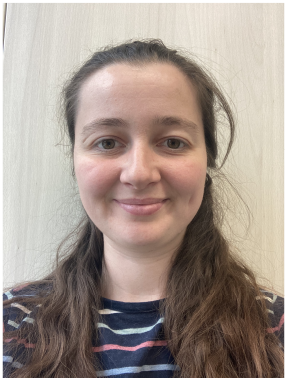

**Freya I. Addison** is a Post-Doc specialising in the development of tools to monitor insect activity using weather instruments. She did her PhD at the University of Leeds in the United Kingdom under the Natural Environment and Research Council Doctoral Training Partnership. Her background is in physics, and previous work includes the development of methodology for simulating the radar cross sections of insects.

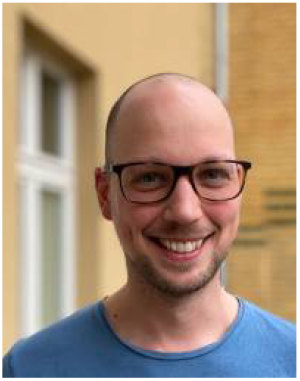

**Maximilian Maahn** received the M.S. degree in meteorology from the University of Bonn, Bonn, Germany, in 2010, and the Ph.D. degree in meteorology from the University of Cologne, Cologne, Germany, in 2015. He was the Research Scientist of the University of Cologne from 2011 to 2016, after which he moved to the Cooperative Institute for Research in Environmental Sciences (CIRES) of the NOAA Earth System Research Laboratories and the University of Colorado at Boulder, Boulder, CO, USA, from 2016 to 2020. Since 2020, he leads the clouD and pRecipitation Observations for Process Studies (drOPS) Group, Leipzig University, Leipzig, Germany. His main research interests include improvement of remote sensing and in situ observations of clouds and precipitation. He is currently developing a novel in situ sensor for snowfall. Dr. Maahn is a member of the American Meteorological Society and the German Meteorological Society. He is an Associate Editor of the Atmospheric Measurement Techniques journal.

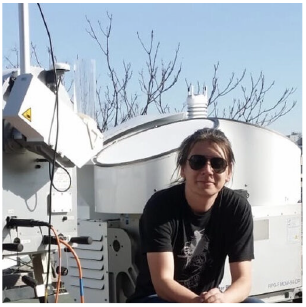

**Willi Schimmel** is a Post-Doc specializing in atmospheric dynamics and modeling. He completed his degree in Applied Mathematics at Leipzig University of Applied Sciences. His research focused on the numerical simulation of large chemical multiphase systems in the troposphere. Schimmel earned his Ph.D. at the Leipzig Institute for Meteorology, where he concentrated on cloud droplet identification using machine learning methods. Currently, Schimmel’s research revolves around Polarimetric Radar Signatures of Ice Formation Pathways from Controlled Aerosol Perturbations, combining cloud seeding, remote-sensing observations, and numerical modeling. His work will contribute to advancing our understanding of atmospheric processes, i.e. ice crystal formation and their impacts.

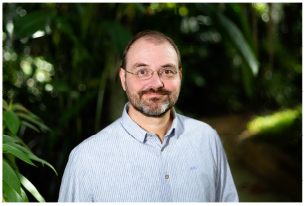

**Christian Wirth** is a plant ecologist who studies how natural and human-induced changes in plant biodiversity affect ecosystem processes. Prof. Wirth leads the working group for Special Botany and Functional Biodiversity at Leipzig University together with Prof. Alexandra Weigelt. He is the founding director of the DFG Center “German Center for Integrative Biodiversity Research (iDiv) Halle-Jena-Leipzig”, director of the Botanical Garden of Leipzig University and associate member of the Max-Planck-Institute for Biogeochemistry in Jena from 2012-2022. 2024 he was admitted to the Leopoldina and to the Saxon Academy of Sciences and Humanities. He received his PhD in 2000 for his work linking carbon cycle and fire ecology of boreal forests in Siberia. Subsequently - with stations in Princeton, Fairbanks and Jena - he developed the TRY database and trait-based approaches to study the relationship between biodiversity and ecosystem functions in forest and grassland ecosystems. More recently, he has been conducting research on forest conservation, floodplain ecology, canopy ecology (Leipzig Canopy Crane), and the impact of climate change on forest biodiversity and ecosystem services.

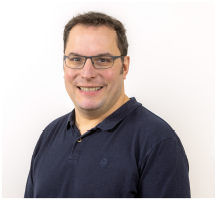

**Johannes Quaas** is professor for Theoretical Meteorology at Leipzig University. He specialises in clouds and climate, with expertise in atmospheric modelling and satellite remote sensing, as well as in biodiversity - climate interactions. Quaas was lead author for the recent assessment report by the Intergovernmental Panel on Climate Change.

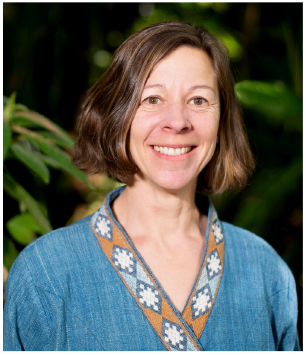

**Heike Kalesse-Los** received her Ph.D. degree in meteorology from Johannes Gutenberg University, Mainz, Germany, in 2010. She was a Post-Doctoral Researcher with McGill University, Montreal, QC, Canada, from 2011 to 2014. She is Professor of Arctic climate change and remote sensing of the atmosphere at Leipzig University, Leipzig, Germany. Her research centers on microphysical process understanding of clouds and precipitation formation around the globe with a focus on Arctic mixed phase clouds, Southern Ocean clouds, and trade-wind clouds. In her work, she combines synergistic profiling of the atmosphere with active and passive remote sensing instruments with machine-learning methods to, e.g., exploit cloud radar Doppler spectra information content for liquid detection and riming occurrence. Recently, she has also become interested in using atmospheric remote sensing observations for novel deployments, e.g., in the field of interactions between climate change and biodiversity change research.

## Notes

### Competing Interest Statement

The authors have declared no competing interest.

### Summary of Updates

The underlying data set changed to facilitate a better comparison between aerial insect concentrations derived from Doppler cloud radar observations from different frequencies.

